# Calcium modulates intramolecular long-range contacts to form a polymorphic α-synuclein A53T fibril

**DOI:** 10.64898/2026.03.14.711779

**Authors:** Jessica Y.C. Huang, Kuen-Phon Wu

**Affiliations:** Institute of Biological Chemistry, Academia Sinica, Taipei, Taiwan; Institute of Biochemical Sciences, National Taiwan University, Taipei, Taiwan

**Keywords:** α-synuclein, amyloid fibril, paramagnetic relaxation enhancement, calcium, long-range contacts, cryo-EM

## Abstract

Human α-synuclein (aSyn) is an intrinsically disordered protein, and aggregations of its amyloid fibrils are associated with Parkinson’s disease (PD). Apart from familial aSyn mutations, accumulated environmental calcium exacerbates aSyn aggregation and accelerates symptoms in aging PD patients. Here, we explored the effects of Ca^2+^ ions on aSyn A53T, an aggregation-prone mutant variant, from disordered states to fibrillar structures. Paramagnetic nuclear magnetic resonance (NMR) revealed that binding of Ca^2+^ ions to the aSyn C-terminal (residues 110-140) relaxed the aSyn conformation, resulting in more aggressive fibrillogenesis. Cryo-electron microscopy structures of aSyn A53T with or without Ca^2+^ ion revealed substantial differences in amyloid folds and fibril assemblies. We characterized N1 (residues 61-66), N2 (residues 69-79), and N3 (residues 89-95) segments in the central non-amyloid β component (NAC) crucial for forming localized structural contacts during early-step aggregation. Our work establishes the contacts governing aSyn misfolding from disordered monomer to aggregated fibril and provides insights into the structural changes elicited by Ca^2+^ ions.

## Introduction

Alpha-synuclein (aSyn) is an intrinsically disordered protein (IDP)^1^ lacking a static tertiary structure and that is primarily located at the termini of presynaptic nerves^2^. Progressive conversion of disordered aSyn from a monomeric state to fibrillar aggregates that are deposited in neuronal cells is a hallmark of Parkinson’s disease (PD), dementia with Lewy bodies, and multiple system atrophy (MSA)^3, 4^. More than 10 million people worldwide currently suffer from neurodegenerative PD and related synucleinopathies^5^. Many factors are known to underlie aSyn conformational changes, including 10 hereditary mutations (Fig. 1A)^6–13^, post-translational modifications (PTMs)^14, 15^, environmental pH, and divalent metal ions^16^, hence contributing to varying rates of fibrillar formation^5^. As yet, the stepwise mechanism by which aSyn is converted from a monomeric IDP to a well-ordered fibril has not been fully elucidated. The polymorphism of aSyn fibrils visualized by cryo-EM has indicated that the protein misfolding issue is even more complex^17–22^.

**Figure 1.**
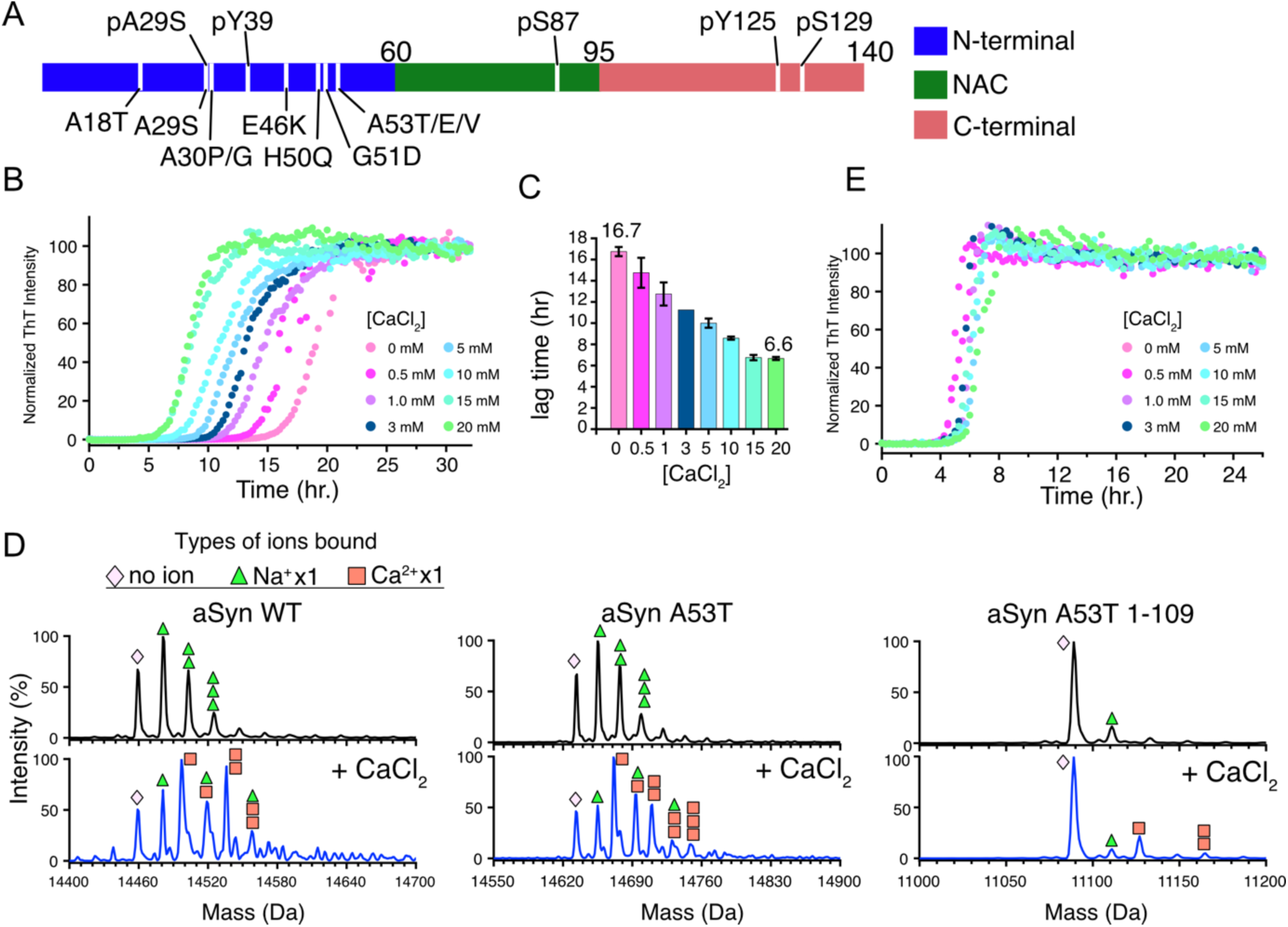
Calcium primarily binds to aSyn at its C-terminal region. (A). Division of the aSyn sequence into three regions: N-terminal (blue, residues 1-60), NAC (green, residues 61-95), and C-terminal (red, residues 96-140). To date, ten aSyn mutations and five post-translational modifications (phosphorylation) have been reported, each of which influences aSyn aggregation and fibril formation, and the positions of these are labeled. (B) ThT time-course aggregation kinetics for aSyn A53T in the presence of CaCl_2_ (0 to 20 mM). Data-points represent averages from triplicate experiments and have been normalized to facilitate comparisons. (C). The lag time of aSyn A53T aggregation kinetics, as shown in (B), can be dramatically reduced from 6.7 to 6.6 h by increasing the [CaCl_2_] from 0 to 20 mM, respectively. (D). Electrospray ionization mass spectrometry (ESI-MS) was used to verify the specificity of aSyn-Ca^2+^ binding. Three aSyn variants (left: wild type, middle: aSyn A53T, and right: C-terminal-truncated aSyn A53T (residues 1-109)) were mixed with 0 or 3.6 mM CaCl_2_. Mass values in the presence of CaCl_2_ indicate a considerable proportion of both wild type aSyn and aSyn A53T bind to Ca^2+^. The pink diamond, green triangle, and orange box symbols represent the mass of native aSyn, and addition of the mass of a single Na^+^ or Ca^2+^ ion on aSyn, respectively. C-terminal-truncated aSyn A53T lacks signal for calcium-bound species, indicating that the C-terminal region (110-140) is the primary Ca^2+^ binding site. (E). ThT aggregation kinetics of C-terminal-truncated aSyn A53T with different [CaCl_2_] confirms the ESI-MS data showing that Ca^2+^ binds to the C-terminal region and accelerates aSyn aggregation since aggregation kinetics are only mildly affected by altered Ca^2+^ concentrations in the absence of the C-terminal (residues 110-140).

aSyn is a 140-residue protein comprised of three defined regions: an amphipathic N-terminal (residues 1-60), a central NAC (residues 61-95), and an acidic C-terminal (residues 96-140). Although aSyn is intrinsically disordered, several intramolecular long-range contacts exist between the N- and C-termini, enabling heterogeneous monomeric assemblies of aSyn, as characterized by solution NMR spectroscopy^23–26^ and single-particle fluorescence spectroscopy^27^. Previous studies have shown that the strengths of these long-range contacts are greatly reduced by the single A53T point mutation^28^. Thus, aSyn A53T fibrils form more readily than wild type aSyn, with the former being linked to early-onset PD^6^. In contrast to the diverse ensembles of disordered aSyn, aSyn fibrils all appear to share a common kernel, with residues 35-100 having been characterized as localized in the amyloid core of the protein^4^. It is worth noting that the N- and C-termini are not directly folded in the aSyn fibrillar structures.

Apart from mutations, divalent metal ions (i.e., Ca^2+^, Mg^2+^, and Cu(I or II)) have been found to interact with aSyn and to increase aSyn fibrillization rates *in vitro*^16, 29^. For instance, Cu(I) induced formation of a local helical structure at N-terminal residues 1-15 of aSyn and disrupted localized binding of neighboring residues 50 and 116-127^30^. Ca^2+^ ion is a secondary messenger responsible for many physiological functions, including neurotransmitter release. Ca^2+^ homeostasis within neurons is vital to maintain cellular health and function. A 35% increase in resting Ca^2+^ ion levels has been observed in the neurites of Alzheimer’s patients, with calcium dyshomeostasis being associated with formation of Lewy bodies in those patients^31^. Calcium homeostasis is regulated via complex interplay between ion channels and cellular organelles (endoplasmic reticulum, mitochondria, and lysosomes), and it may become defective with aging^32^. If calcium homeostasis is disrupted, the accumulated intracellular Ca^2+^ ions can drive aSyn oligomerization and aSyn fibrillization, ultimately causing neuronal defects^33^. However, as yet, a structural analysis of aSyn fibril formation in the presence of calcium has not been reported.

In this study, we adopted a multidisciplinary biophysical approach encompassing Thioflavin T (ThT) aggregation kinetics, mass spectrometry, solution NMR spectroscopy, and cryo-EM to reveal how Ca^2+^ ion alters aSyn assemblies and fibrillogenesis. We chose to focus on the familial aSyn A53T mutant, revealing a combinatorial effect of the mutant variant and Ca^2+^ presence on aSyn aggregation kinetics. Our mass spectrometry and NMR analyses confirm that aSyn A53T hosts calcium binding sites in its C-terminus. Paramagnetic relaxation enhancement (PRE) NMR further demonstrate that long-range contacts are substantially reduced in the presence of Ca^2+^ ions, resulting in a relaxed ‘open’ conformation that enables rapid aggregation kinetics. Moreover, cryo-EM revealed distinctive aSyn A53T fibrillar structures in the presence or absence of Ca^2+^ ions. Our findings demonstrate that calcium drives conformational changes in aSyn A53T, switching it from a disordered resting state to polymorphic fibrils.

## Results

### Calcium ions bind to the C-terminal region of aSyn and accelerate fibril formation

We measured the aggregation kinetics of aSyn A53T in the presence of varied CaCl_2_ concentrations. Thioflavin T (ThT) kinetic analysis revealed rapid aSyn A53T fibril formation when 20 mM CaCl_2_ was added to solution (Fig. 1B), consistent with previous reports^29, 30, 34^. The lag time for aSyn A53T fibril formation decreased from 16.7 h without CaCl_2_ to <7 hours when the CaCl_2_ concentration was >15 mM (Fig. 1C). The maturation time of aSyn A53T fibrils was also reduced from ∼30 h (without CaCl_2_) to 10 h in the presence of 20 mM CaCl_2_. The ∼60% reduction in aggregation lag time and maturation time support that calcium ions play a critical role in accelerating aSyn A53T fibrillogenesis. We detected similar effects for MgCl_2_ (Supplementary Fig. 1), with divalent Mg^2+^ also significantly promoting aSyn A53T fibril formation. Thus, our ThT kinetics analysis indicates that the aggregation-prone aSyn A53T variant is more reactive in terms of forming fibrils when divalent metal ions are present.

The C-terminal region of aSyn has been postulated as the primary region responsible for calcium-binding^34, 35^. We used electrospray ionization (ESI) mass spectroscopy to measure changes in mass of wild type aSyn, aSyn A53T, and C-terminal-truncated aSyn A53T ΔC (residues 110-140 deleted) in the absence or presence of CaCl_2_ (Fig. 1D). We detected multiple Ca^2+^-bound states (1-3 Ca^2+^) for the protein’s C-terminal region, as reported previously^36–38^. As anticipated, there was no significant population of calcium-bound aSyn A53T ΔC in the ESI MS spectra. Moreover, the ThT kinetics of aSyn A53T ΔC (Fig. 1E) across a broad range of CaCl_2_ concentrations (0 - 20 mM) were equivalent. These results demonstrate that aSyn responds to environmental ions mainly through its negatively-charged C-terminal region.

### NMR characterization of calcium-bound residues and protein conformation

To investigate the impact of CaCl_2_ on aSyn A53T fibrillar formation, we determined if monomeric aSyn A53T conformation was dramatically altered by Ca^2+^. NMR-monitored titration experiments of aSyn A53T with CaCl_2_ (0-20 mM) (Fig. 2A, Supplementary Fig. 2) revealed that residues in the C-terminus, as well as in the N-terminal and NAC regions, were disrupted by calcium in solution. Nevertheless, the disordered state of aSyn A53T did not appear to change in the presence or absence of CaCl_2_, based on the NMR spectra. However, negatively-charged residues—including E83, E104, D115, D119, D121, and E130—did appear to be perturbed, and residues K21, T53, K58, T59, T92, and S129 proved sensitive to the presence of environmental calcium. Thus, the hydroxyl groups of serine and threonine residues in the A53T mutant may play an important role in binding calcium ions. Positively-charged residues K21 and K48 follow precede glutamine and threonine residues, respectively, potentially serving to bind calcium ions and alter the local chemical environment^35^.

**Figure 2.**
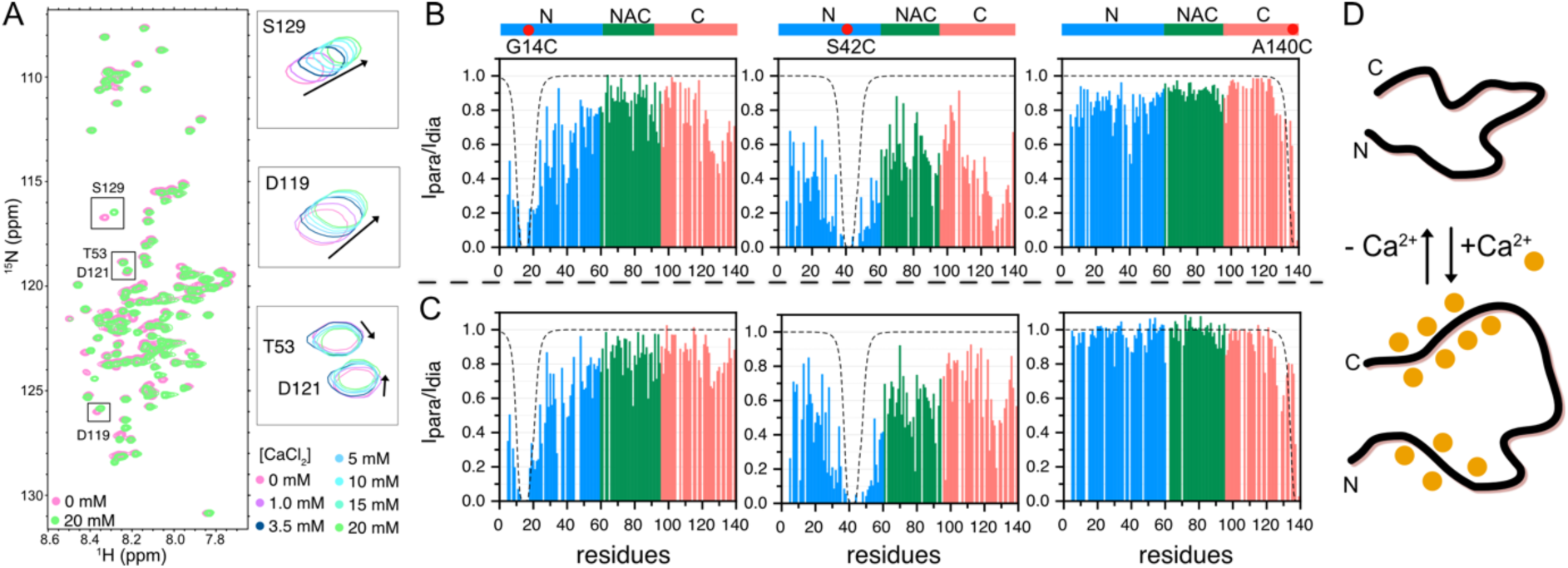
NMR characterization of the tertiary interactions of Ca^2+^-bound aSyn A53T. (A). aSyn A53T was titrated with 0-20 mM CaCl_2_ and assessed by means of NMR ^15^N-HSQC spectra. The C-terminal region is significantly perturbed under conditions of low [CaCl_2_] (i.e., 1 mM) (Supplementary Fig. 2). However, in the presence of 20 mM CaCl_2_, many residues across the entire aSyn A53T sequence are affected by Ca^2+^ (Supplementary Fig. 2). Examples of cross-peak shifts for residues S129, D119, D121, and T53 are shown. Results from paramagnetic relaxation enhancement NMR of aSyn A53T in the absence or presence of 20 mM CaCl_2_ are shown in (B) and (C), respectively. Three separate plots are shown to illustrate the PRE effects of the G14C, S42C, and A140C spin labels (red dots), respectively. The dashed curves in each plot demonstrate the PRE effect on a fully linearized 140-aa protein^74^. The N-terminal, NAC, and C-terminal regions are colored blue, green, and salmon in each plot. (D). Schematics of aSyn A53T showing the conformational changes upon addition or removal of Ca^2+^ ions (orange dots).

Next, we evaluated changes in the tertiary conformation of aSyn A53T upon adding 20 mM CaCl_2_. PRE-NMR is a sensitive tool for detecting changes in long-range contacts (i.e., up to 25 Å) of disordered synuclein^24, 26, 39^ at the resolution of single residues. For our PRE-NMR experiments, we separately mutated three aSyn residues (G14, S42, and A140) to cysteine to conjugate the spin label MTSL (Fig. 2B, 2C). The aSyn A53T G14C-MTSL construct allowed us to detect long-range interactions between N-terminal G14C and essential residues 124-140 in the C-terminal region. The reverse spin label on A140C consistently revealed that the N- and C-terminal regions of aSyn exhibit long-range interactions, as NMR cross-peak signals were reduced for the N-terminal residues. NMR signal for the S42C spin label was globally reduced, particularly for neighboring residues (amino acids 30-60) and C-terminal residues 125-140, demonstrating that S42 and its neighbors are pivotal in maintaining the tertiary conformational contacts of aSyn monomers.

Notably, the intramolecular long-range contacts between the N- and C-terminal regions were significantly reduced when aSyn A53T spin label samples were mixed with CaCl_2_ (Fig. 2C). For instance, the relative intensity peak ratio (I_para_/I_dia_) of C-terminal residues 125-140 in the presence of CaCl_2_ is ∼0.8 for the G14C spin label compared to 0.45-0.6 without CaCl_2_, indicative of an extensive distance between the spin label probe and the C-terminal. Similarly, the A140C-MTSL construct in the presence of CaCl_2_ exhibited nearly no contacts from residue 140 to the N-terminal or NAC regions of aSyn A53T. The S42C spin label induced significantly reduced paramagnetic relaxation effects when calcium ions were bound to aSyn A53T. The limited long-range interactions of aSyn A53T around residues 125-140 in CaCl_2_ solution indicates that the calcium-bound C-terminal region of aSyn A53T is isolated in the solvent, resulting in its reduced interactions with the first 100 residues of the synuclein protein. Intriguingly, a P1 motif that hosts the S42 residue was proposed previously as being critical to aSyn fibril formation^39^. The reduction in long-range contacts when calcium ions are present evidenced by our PRE data on residue S42 confirms the importance of this P1 motif to synuclein fibrillogenesis and its sensitivity to environmental factors. Our PRE-NMR results illustrating increased radial size for a disordered ensemble conformation of aSyn A53T monomer in the presence of Ca^2+^ ions (Fig. 2D) supports a previous small-angle X-ray scattering (SAXS) study^37^ that reported a ∼10% increase in the radius of gyration of aSyn when mixed with caclium.

### Cryo-EM structures of polymorphic aSyn A53T fibrils

The distinct impact of calcium on the conformation of monomeric ensembles of mutant synuclein and their corresponding aggregation kinetics prompted us to investigate if aSyn A53T forms consistent fibril structures. A cryo-EM analysis previously revealed polymorphism among more than two dozen synuclein fibrils^17^ under various conditions, but the structure of Ca^2+^- or metal ion-induced aSyn fibrils had remained utterly unknown. Thus, we collected two cryo-EM datasets; one for aSyn A53T fibrils incubated with CaCl_2_ and one without. The cryo-EM map of aSyn A53T alone (termed aSyn A53T hereafter) has an overall resolution of 3.4 Å, and the map of aSyn A53T fibrils formed in the presence of CaCl_2_ (termed aSyn A53T-Ca) was determined at 2.7 Å (Fig. 3 and Supplementary Fig. 3). The aSyn A53T structure comprises two protofilaments (PFs) with a crossover distance of ∼600 Å, in which ten β sheets in the amyloid core of an individual aSyn A53T chain (residues 36-100) could be defined (Fig. 3A-C, 3G). The two PFs of aSyn A53T assemble via a compact staggered steric zipping mechanism, with ^50^HGVTTAVE^57^ forming the interface (Fig. 3B, 3C). In our aSyn A53T fibril structure, residue 53 faces the PF interface and is directly involved in fibril assembly. That structure exhibits the same fold as a previously determined aSyn A53T fibril (PDB ID: 6LRQ)^40^, but the assembly point and interface are completely different, most notably in terms of A53T being directly involved in PF assembly in our structure. In the previously reported aSyn A53T fibril structure^40^, fibril assembly is mediated through residues T59 and L60, and T53 is not involved. Intriguingly, our aSyn A53T fibril structure exhibits both the same fold and assembly pattern as a C-terminal-truncated aSyn fibril (residues 1-121) (PDB ID: 6H6B)^41^. Removing the C-terminal region of aSyn has been found to accelerate aSyn aggregation^42^, as confirmed by our ThT kinetic assays (Fig. 1E). The matching fibrillar structures of C-terminal-truncated aSyn and the aggregation-prone A53T variant confirm that intramolecular interactions from the C-terminal region to the NAC and N-terminal regions are crucial to prevent monomeric aSyn assemblies from rapidly aggregating^26, 28, 41^.

**Figure 3.**
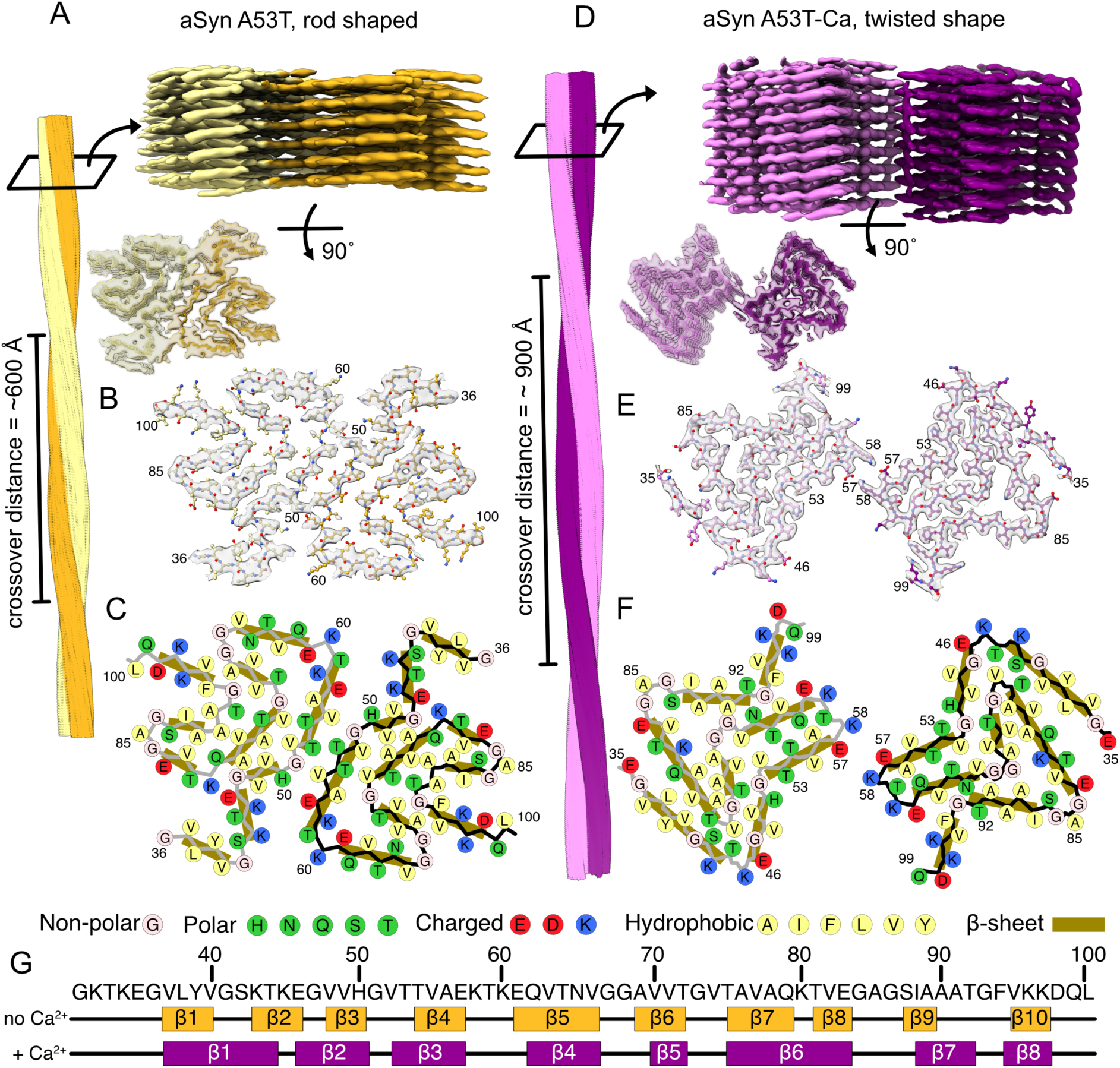
Cryo-EM structures of aSyn A53T and Ca^2+^-bound aSyn A53T fibrils. (A). aSyn A53T fibril in which two PFs (yellow and gold) are assembled into a rod-shaped filament with a crossover distance of ∼600 Å. (B). Cross-section of the aSyn A53T fibril showing two aSyn A53T chains. (C). Detail of the aSyn A53T cross-section depicting residues and β-sheets and showing how residues H50-E57 closely tether the two PFs. (D). The aSyn A53T-Ca fibril of two PFs (pink and violet) exhibits a large crossover distance of ∼900 Å and appears twisted. (E). Cross-section of the aSyn A53T-Ca fibril. (F). As for (C), but the assembled aSyn A53T-Ca fibril shows how charge-charge interface interactions alone drive fibril assembly. (G). The aSyn A53T and aSyn A53T-Ca fibril structures have 10 and 8 β-sheets, respectively. Two long β-sheets in aSyn A53T-Ca resulted in it having fewer sheets.

Our aSyn A53T-Ca fibril structure presents a twisted shape with a ∼900 Å crossover distance, i.e., considerably larger than for the aSyn A53T fibril in the absence of calcium (Fig. 3D). We characterized eight β sheets between residues 35-99 from our 2.7 Å map of aSyn A53T-Ca fibril (Fig. 3G). The aSyn A53T-Ca fibril comprises two PFs, with the assembly being stabilized by recurring hydrogen-bonding between aSyn fibril layers. Folding of aSyn A53T-Ca is notably different from aSyn A53T described above, but is similar to PDB ID 6SSX^43^ and patient fibril-derived structures (PDB ID codes 7NCA, 7NCG, 7NCH, 7NCI, and 7NCJ^44^, which have been described as having a “sandal-like” fold, as opposed to the “boot-like” fold of aSyn A53T). Nevertheless, our aSyn A53T-Ca fibril and these other aSyn fibril structures are assembled distinctively. As shown in Figure 3F, the two charged residues K57 and E58 represent the interface between the two PFs of aSyn A53T-Ca, with inter-PF E57-K58 sidechain hydrogen bonds (H-bonds) tightly linking the two PFs. The point-to-point assembly has been reported previously for aSyn fibrils, including for Parkinson’s disease-related Y39-phosphorylated aSyn (6L1T)^45^ and aSyn fibrils seeded from MSA patients (7NCI, 7NCJ)^44^. Nevertheless, the aSyn A53T-Ca fibril structure we present here represents a new polymorph. Surprisingly, the mutated T53 residue of this aSyn A53T-Ca fibril structure is solvent-exposed and does not interact with the other PF.

### Structural details of aSyn A53T and aSyn A53T-Ca fibrils

The two aSyn fibril structures we have determined herein (aSyn A53T and aSyn A53T-Ca) exhibit completely different folding patterns and assemblies, clearly reflecting their respective compact or relaxed monomeric conformations and aggregation kinetics. Inspecting the intrachain folds and their H-bonds, hydrophobic patches, and surface electrostatic charges provided insights into the mechanism of rapid aSyn aggregation. The boot-shaped fold of aSyn A53T is primarily restricted to residues 36-100 in our cryo-EM fibril structure (Fig. 4A, Supplementary Fig. S4A-C). Its V49, V52, V74, A76, V77, A78, I88, A90, A90, A91, and F94 hydrophobic residues together constitute a massive hydrophobic patch across the boot-shaped fold (Fig. 4A). In addition to H-bonds formed across β sheets, we identified four intramolecular H-bonds within the boot-shaped fold involving residue pairs Y39-E46, K58-E61, G73-T75, and E83/G84-S87 (Fig. 5A). None of these four H-bonds is associated with long-range bonding, implying that aSyn A53T folding is essentially maintained by hydrophobicity.

**Figure 4.**
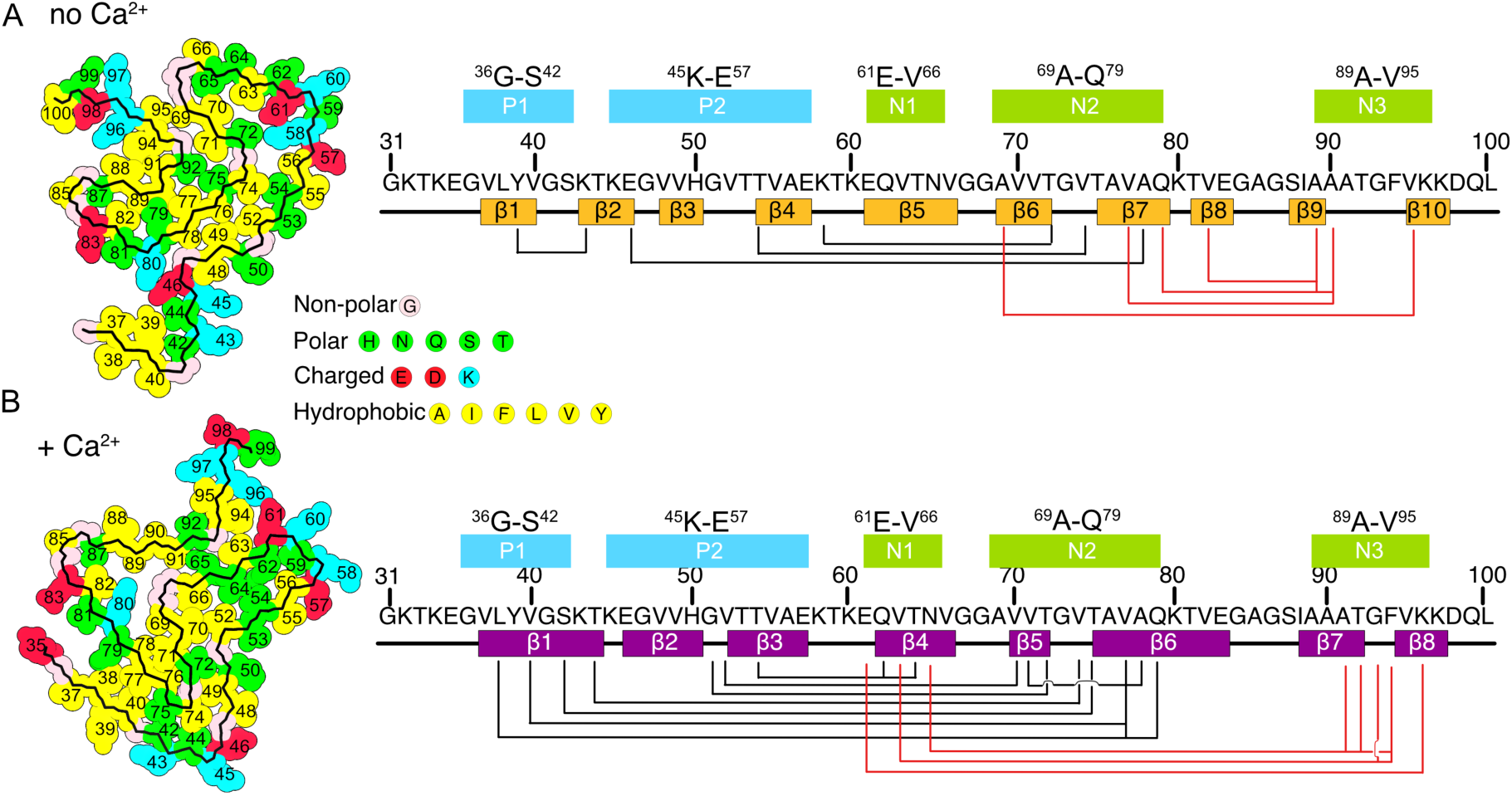
Long-range intrachain interactions determine the distinct folds of aSyn A53T and aSyn A53T-Ca fibrils. Many interactions within the aSyn A53T (A) and aSyn A53T-Ca (B) structures maintain their folded amyloid cores. Polar, negatively-charged, positively-charged, and hydrophobic residues are colored green, red, blue, and yellow, respectively. Hydrophobic interactions play a critical role in both structures, but aSyn A53T-Ca has a larger hydrophobic patch than aSyn A53T. The van der Waal’s interactions of both amyloid structures were analyzed using ChimeraX 1.3^75^. Long-range contacts (inter-residue distance ≥5 residues) are illustrated, revealing five key regions—the P1 and P2 motifs^39^ and three novel regions N1, N2, and N3 in the NAC identified in our study—responsible for the folding of amyloid cores. Long-range contacts within the NAC are colored red to distinguish them from interactions between the N-terminal region (P1, P2) and the NAC (black).

**Figure 5.**
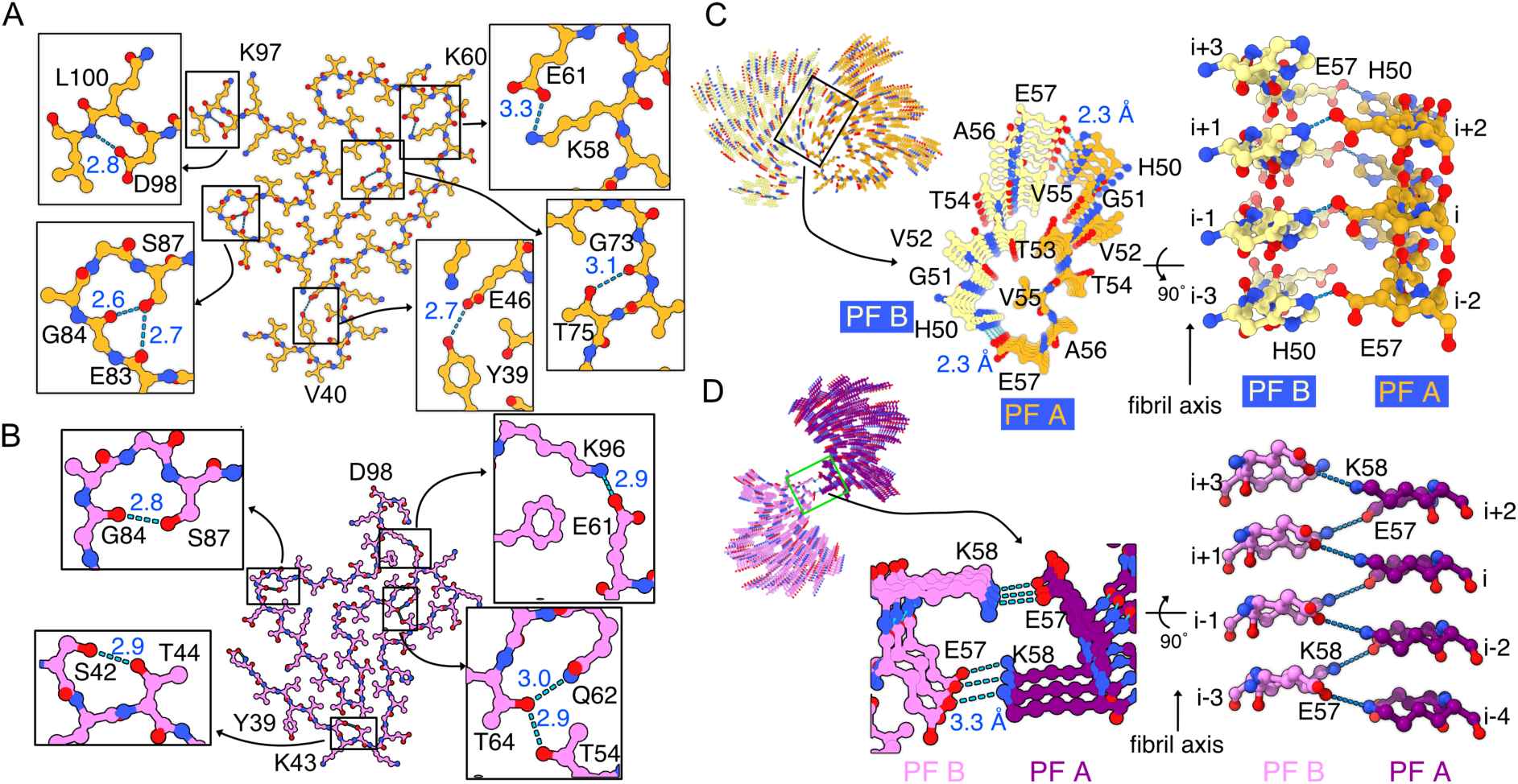
Hydrogen bonding networks of aSyn A53T and aSyn A53T-Ca fibrils. H-bonds in the aSyn A53T and aSyn A53T-Ca fibrils stabilize the folded conformation. (A-B). We identified 5 and 4 primary intrachain H-bonds in aSyn A53T and aSyn A53T-Ca fibrils, respectively. aSyn A53T-Ca has two long-range H-bonds (E61-K96 and T54-T64) that could potently stabilize and accelerate the folding process during oligomerization. (C-D). Both fibril types also have inter-protofilament H-bonds that tightly constrain the fibril structure. The inter-protofilament H-bonds are consistent across three chains of two PFs. In aSyn A53T fibrils, residues H50 and E57 form the inter-protofilament H-bonds, whereas it is E57 and K58 that do so in aSyn A53T-Ca fibrils. All H-bonds are indicated by blue dashed lines and the bond distances are shown.

The structural unit of aSyn A53T-Ca fibrils folds differently from that of aSyn A53T fibrils. Residues 36-99 of the sandal-like aSyn A53T-Ca structure fold into two stacked V-shaped β-arch layers (Fig. 4B, Supplementary Fig. S4A-C), and its 1847 Å^2^ hydrophobic patch is more extensive than that of the aSyn A53T fibril structure (1532 Å^2^). This hydrophobic patch covers the two β-arches (encompassing four layers)—involving residues L38, V40, V49, V52, V66, A69, V70, V71, V74, A76, V77, and A78—in which three hydrophobic long-range clusters play a critical role in maintaining the sandal-like fold. According to the intrachain contacts we have identified from the two aSyn fibril types, we have characterized three segments in the NAC (residues 61-95), denoted N1 (residues ^61^EQVTNV^66^), N2 (^69^AVVTGVTAVAQ^79^), and N3 (^89^AAATGFV^95^), which strongly interact with each other and with previously identified P1 (^36^GVLYVGS^42^) and P2 (^45^KEGVVHGVTTVAE^57^) motifs^39^ to form distinct fibrillar folds. Intrachain S42-T44, T54-Q62/T64, E61-K96, and G84-S87 H-bonds in aSyn A53T-Ca also provide additional stabilization (Fig. 5B). The long-range T54-T64 and E61-K96 H-bonds are fundamental for sustaining the sandal-like fold and the hydrophobicity-driven folding process of the aSyn A53T-Ca fibril structure.

We identified three hydrophobic clusters in aSyn A53T-Ca fibrils. The first hydrophobic cluster consisting of residues L38, V40 and V77 brings the P1 and N2 regions into close proximity, but these two regions do not interact in the aSyn A53T fibril structure. The second cluster properly staggers the valine sidechains and retains contact between the P2 and N1 regions. The third cluster comprising ^63^V-N^65^ and ^91^A-F^94^ ensures close contact between the N1 and N3 regions. Apart from these hydrophobic clusters, intramolecular long-range E61-K96 and T54-T64 H-bonds provide additional force to stabilize the stacked V-shaped fold. Thus, the three hydrophobic clusters and the H-bonds across the amyloid core of the aSyn A53T-Ca structure are crucial for stably preserving synuclein folding during rapid fibrillar extension and assembly (Fig. 4B).

The two aSyn A53T and aSyn A53T-Ca fibril structures we have determined assemble via completely distinct contacts. The interface residues ^50^H-E^57^ (in the P2 region) of aSyn A53T fibrils stagger T53 and V55 spatially, with the H50 and E57 sidechains forming H-bonds (Fig. 5C). Residues E57 and H50 of one PF (i.e., chain i of PF A) separately form H-bonds with H50 of the preceding chain (i-1) and E57 of the next chain (i+1) of the second PF (PF B). The H-bond interchain distance between Nε_2_ of H50 and Oε_2_ of E57 is 2.3 Å, indicative of a compact assembly. In contrast, aSyn A53T-Ca protofilament assembly is simplified by a point-to-point contact. The neighboring oppositely-charged residues E57 and K58 in chain i of PF A form H bonds to K58 in chain i-1 and E57 in chain i+1 of PF B, respectively (Fig. 5D). The interchain H-bond distance between Oε_1_ of E57 and Nσ of K58 is 3.3 Å. It is worth noting that E57 forms H-bonds in both types of fibril assemblies, but the residues it pairs with are not identical. Notably, aSyn A53T and aSyn A53T-Ca fibrils both require hydrogen bonding among the three chain layers of two PFs to lock the assembled fibrils tightly together.

### Water molecules in the aSyn A53T-Ca structure enhance fibril stabilization

There is a large cavity near ^79^Q-A^91^ in the sandal-like aSyn-A53T-Ca structure, and this perhaps corresponds to considerable unresolvable potential in the cryo-EM map and likely represents several solvent molecules (Fig. 6A, 6B). A water channel density was identified previously for the aSyn H50Q structure^46^. Six water molecules could be comfortably placed in the unresolved region of our aSyn-A53T-Ca structure, revealing potential H-bonds between these six molecules and aSyn residues G67, A69, Q79, K80, I89, and A89 (Fig. 6B). In addition, three water molecules were apparent in the unresolved portions of cryo-EM aSyn-A53T-Ca maps neighboring the ^70^VVT^72^ (Fig. 6C) and ^51^GVTT^54^ (Fig. 6D) regions, highlighting interactions between those water molecules and residues G51, T53, T54, V70, and T72. Four of the nine characterized water molecules are located between two stacking aSyn A53T-Ca chains (layers) and form H-bonds between the layers, thereby stabilizing aSyn A53T-Ca during PF oligomerization and extension. Thus, incorporation of water molecules by aSyn-A53T fibrils may provide greater stabilization of the boot-shaped fold and fibril layers. However, the nominal resolution of the aSyn A53T structure is only 3.4 Å, rendering it difficult to clearly identify aSyn-water interactions, hence further structural studies are required.

**Figure 6.**
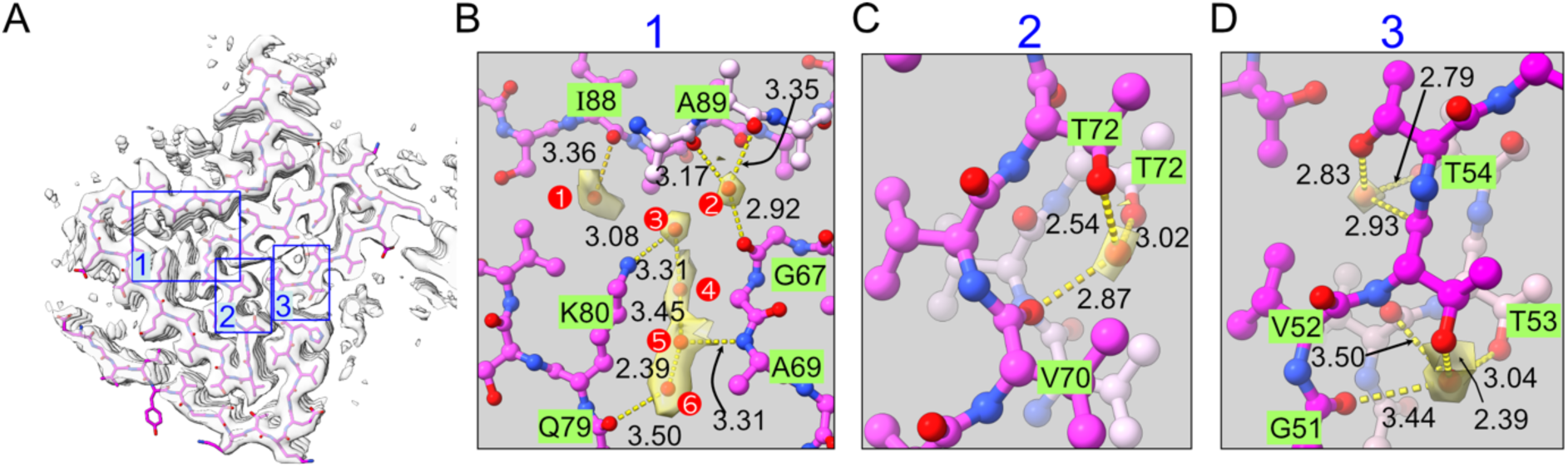
Role of solvent-derived water molecules in the aSyn A53T-Ca structure. (A). Solvent water molecules were identified in the aSyn A53T-Ca map, revealing additional H-bonds that enhance folding during aggregation. Water molecules in (B-D) are shown as red spheres with yellow densities. We identified three regions in the structure map that contain water, but region 1 (B) has the most significant unresolved density representing six water molecules. Those six water molecules can form H-bonds among each other and with aSyn A53T-Ca. Regions 2 (C) and 3 (D) have one and two water molecules, respectively. These water molecules are located between two layers of a single PF and form H-bonds that, in addition to cross-β H-bonds, stabilize the inter-layer formation.

## Discussion

The two cryo-EM fibrillar structures of aSyn A53T we have determined demonstrate that calcium ions significantly alter the intrinsic long-range contacts of disordered aSyn A53T, resulting in two new polymorphs. aSyn A53T is a mutant variant more prone to aggregation than wild type aSyn^6, 47^, with disease-related consequences. Importantly, we show that in the presence of environmental calcium aSyn A53T develops yet another fibrillar structure that exacerbates aggregation. PRE-NMR revealed substantially relaxed long-range contacts between the N- and C-terminal regions of aSyn A53T, as well as P1-N3 interaction, both influenced by calcium presence. The dynamic intramolecular contacts of the N- and C-termini have been proposed to prevent disordered aSyn from attaining the misfolded fibrillar state^23, 26, 29, 42, 48^. We have defined five regions in the amyloid core of aSyn (P1, P2, N1, N2, and N3) to illustrate the unique intramolecular interactions between the boot- and sandal-like synuclein conformations, respectively (Fig. 7). The P1 and P2 motifs in the N-terminal region are deemed essential for aSyn fibril formation^39^. Our PRE-NMR analysis revealed weakened P1-N3 interactions in the presence of Ca^2+^, implying that they act as a secondary protective contact to prevent disordered aSyn from aggregating. In contrast, P1-N3 contact does not exist within the two typical amyloid folds of aSyn, supporting that rearrangement of P1-N3 contacts may trigger and promote aSyn aggregation (Fig. 7, steps i and ii).

**Figure 7.**
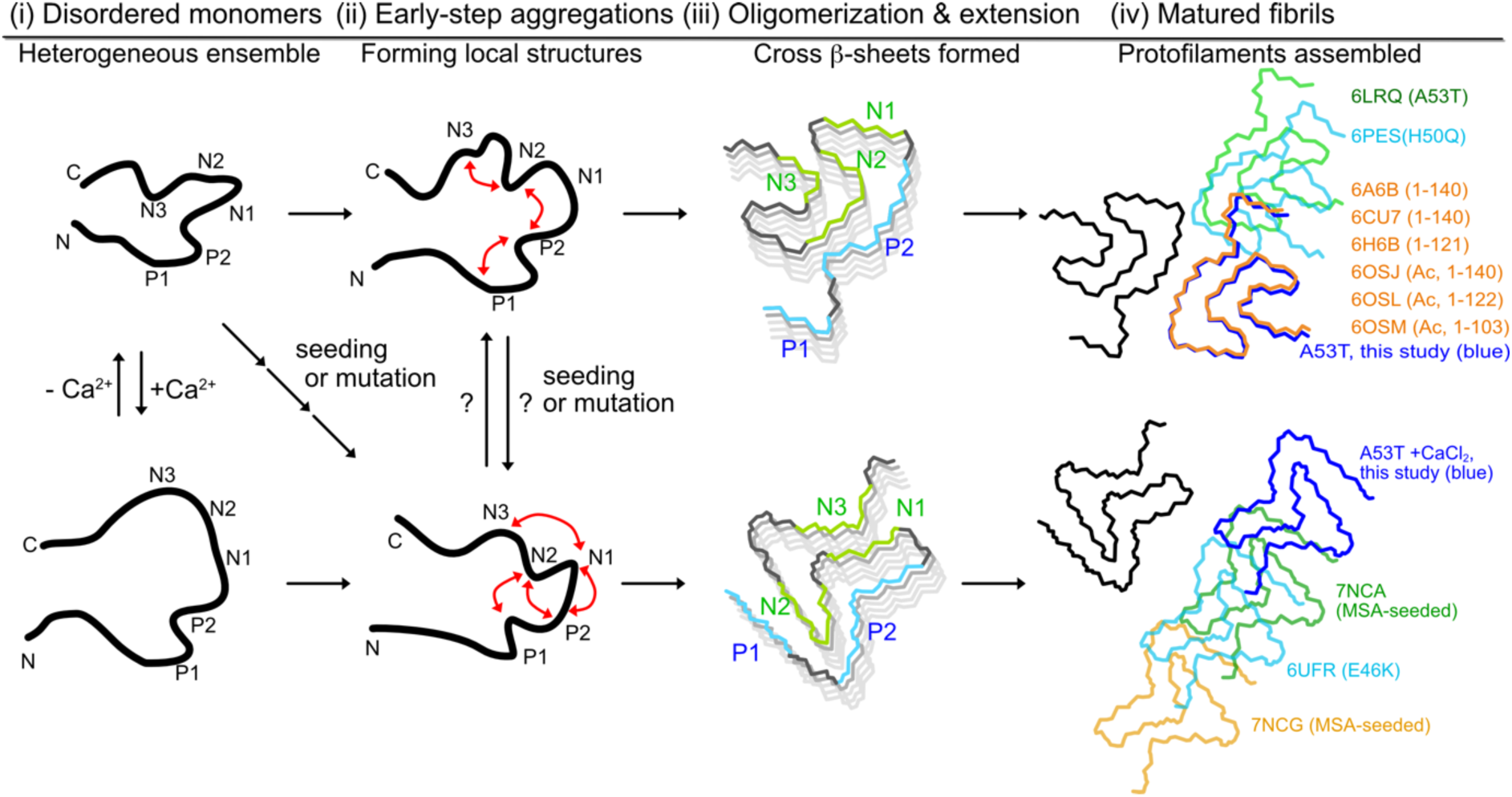
Summary of the aSyn aggregation pathway. The misfolding and aggregation pathway of aSyn is divided into four distinct conformational states. The intrinsically disordered protein ensembles are manipulated by environmental Ca^2+^ (state i), as described in this study, before the distinct formation of intramolecular local contacts (state ii), which precedes oligomerization of the two fibril-like folds (state iii). The two common fold types (“boot-like” or “sandal-like”) of the aSyn amyloid core can be assembled into multiple forms to give rise to polymorphic amyloid fibrils (state iv). The P2 motif is central to facilitating PF contacts. However, the contact points in the boo t-like (top) and sandal-like (bottom) amyloid cores differ. Fibrils in state iv are superimposed for one PF (black), and paired PFs are colored differently to represent polymorphic assemblies. Importantly, sandal-like fibrils primarily assemble via electrostatic attractions combined with inter-protofilament H-bonds.

In the presence of Ca^2+^, equilibrated aSyn monomers are ultimately misfolded into aberrant amyloid folds. It remains unclear if seeding or mutation effectively redistributes protein equilibrium and thus early aggregation steps (Fig. 7, step ii). Once local structural contacts have been established within the protein core during early-step aggregation, it guides aSyn to be oligomerized and extended in the same pattern. Then, interchain cross-β sheet interactions further stabilize the aSyn protofilament (Fig 7. step iii). Recent NMR studies of interactions between aSyn monomer and fibrils indicate that extension of aSyn fibrils is initially mediated by the flanking charged N-terminal residues disordered aSyn^49, 50^. However, it is eminently challenging to capture conformational changes in such heterogeneous and dynamic folding reactions, thus rendering it problematic to characterize aSyn oligomerization entirely. Here, we propose that the defined P1, P2, N1, N2, and N3 regions participate in various interactions underlying aSyn fibrillar formation, resulting in two common aSyn amyloid structures, termed the “boot-like” and “sandal-like” folds. The aSyn A53T-Ca amyloid core exhibits five significant long-range inter-region contacts, including P1-N1, P1-N2, P2-N1, P2-N2 and N1-N3 (Fig. 4B, 7), as well as solvent-aSyn H-bonds (Fig. 6), which result in a large hydrophobic patch and a stable sandal-like fold. The aggregation-promoting P2-N1 and P2-N2 tertiary contacts have been characterized previously in a monomeric aSyn assembly^51^, implying that a mixture of fibril-like and nonfibril-like aSyn can form in solution and that Ca^2+^ ion is responsible for differentiating the aggregated and monomeric aSyn ensembles. In contrast, the mutant aSyn A53T core is maintained by fewer long-range contacts driven by the P2-N2 and N2-N3 networks. Instead, the primary intramolecular contacts of the aSyn A53T core are formed by residues within five amino acids of each other on the protein sequence, resulting in the characteristic boot-like fold.

Although ∼77% (20 of 26) of currently determined aSyn fibrils exhibit either the boot-like or sandal-like conformation in one PF, synuclein PFs still exhibit substantially polymorphic fibrils (Fig 7. step iv). Our aSyn A53T fibrils assembled like other aSyn fibrils, including full-length aSyn (PDB ID: 6A6B^52^ and 6CU7^53^), C-terminal-truncated aSyn (6H6B^40^), and aSyn with an acetylated N-terminus (6OSJ, 6OSL and 6OSM^48^) in which residues 50-59 (the P2 region) act as the interface for sterically zipping together two PFs. In contrast, a limited contact point at residues ^58^KTKE^61^ mediates pairing between two PFs of the hereditary mutant aSyn variants A53T (6LRQ) and H50Q (6PES^46^). Calcium-impacted aSyn A53T-Ca exhibits a folding pattern almost identical to the E46K mutant variant (6UFR^54^) and two MSA-seeded fibrils (7NCA and 7NCG^44^). Nonetheless, all four of these aSyn fibril types generate completely different assembled structures, yet all follow the same rule of electrostatic attraction to tightly link two PFs. In the resulting sandal-like fold of the amyloid core, charged residues K45, E46, E57, and K58 act as the connections. Our aSyn A53T-Ca cryo-EM structure reveals H-bonds between E57 and K58 of two PFs. Similarly, the MSA-seeded fibrils 7NCA and 7NCG utilize E46/K58 or K45/E46 to pair with another PF^44^. The E46K mutant (6UFR^54^) fibril assembles by means of hydrogen bonding between the K45 and E46 residues of two PFs. We note that the hydrophobic and polar residues in the P2 region are not directly involved in PF contacts to generate the sandal-like folded structure, including for the aSyn A53T mutant variant. The surface electrostatic potentials we have generated of the boot-like and sandal-like aSyn amyloid structures depict the pairing mechanism as operating through charge-charge interactions (Supplementary Fig. S4C). Charge-charge-mediated assemblies have been characterized previously for a wide variety of protein fibrils, such as E196K prion^55^, Aβ42^56^, 4R Tau (6TJX^57^), and serum amyloid A (6DSO^58^), indicative of a widespread protein misfolding and aggregation mechanism^59^.

Calcium is an important small molecule associated with neurotransmitter signaling. Our MS, ThT kinetics, and NMR titration data (Fig. 1D, 1E, 2A) conclusively demonstrate that the C-terminal region of the aSyn protein sequence is the calcium-responsive site, consistent with previous ^13^C-NMR and MS studies^35, 38^. We have demonstrated that aSyn A53T fibril structures can be easily manipulated by the presence of environmental Ca^2+^ ions, unlike for other studied hereditary aSyn mutant variants (H50Q, A53T, E46K, G51D)^40, 46, 52, 54, 60^ and aSyn with PTMs^45, 48^. aSyn sequence mutations and posttranslational modifications both result in intrinsic changes to charge distributions and local structural formations, potentially generating different fibrillar structures. However, environmental Ca^2+^ ions can indirectly modify negatively-charged sidechains, primarily in the C-terminal region, thereby reordering aSyn monomer assembly and accelerating aggregation kinetics. Cytosolic Ca^2+^ is normally maintained at a concentration of ∼100 nM, but it may be elevated dramatically (by up to 5000-fold) under conditions of lysosome dysfunction, with these organelles storing 0.4-0.6 mM free calcium ^61^. An imbalance in cytosolic [Ca^2+^] in neuronal cells has been identified as a pathogenic cause of various neurodegenerative diseases^62^. Our study reveals mechanistic details of how aSyn aggregation is impacted by Ca^2+^ ions. Importantly, various divalent metal ions may impact aSyn monomer ensembles differentially, potentially resulting in diverse aggregation pathways. For instance, the Cu^2+^ ion binding site of aSyn is located in the N-terminal region (first 15 residues), with binding of this metal ion resulting in a local helical structure^30^. Combining our aSyn A53T-Ca fibril structure generated herein with further structural studies of aSyn fibrils in the presence of Mg^2+^, Cu^2+^, or Zn^2+^ ions could resolve the misfolding and aggregation pathway of aSyn and clarify the roles of metals in neurodegenerative disorders. Understanding the physiological functions of metal ions in association with aSyn and their pathological outcomes may aid in the development of new therapeutics for Parkinson’s disease.

## Materials and methods

### Protein expression and purification

The full-length aSyn A53T gene was synthesized (Genscript) and subcloned into a pRSFDuet-1 vector with a tag comprising a hexahistidine and TEV protease cleavage sequence (His_6_-TEV-tag). Mutation and truncated synuclein were generated by site-directed mutagenesis and conventional PCR amplification, respectively. All synuclein variants were expressed in *Escherichia coli* (*E. coli*) RIL strain (Novagen). Bacteria were cultured in Luria broth (LB) medium and induced with 0.6 mM IPTG at an OD_600_ of 0.8 for 16 h at 25 _°_C. Cells were pelleted by centrifugation at 6000 rpm, 4 _°_C for 30 min. *E. coli* pellets were resuspended in lysis buffer (25 mM Tris pH 7.6, 200 mM NaCl, 3 mM β-mercaptoethanol (βME), and 0.3 % phenylmethylsulfonyl fluoride) and sonicated. The sonicated *E. coli* were centrifuged at 20,000 rpm, 4 _°_C for 1 h. The synuclein-containing supernatant was processed through a Roche cOmplete Ni-affinity gravity column. Synuclein was eluted with 300 mM imidazole at pH 8.0, before being subjected to TEV protease cleavage and dialysis overnight at 4 _°_C to remove the His_6_-TEV-tag. The dialyzed product was again purified in a Ni-affinity gravity column to trap the His-tagged TEV protease and remove any remaining His_6_-TEV tag. The flow-through and wash fractions were loaded onto a HiTrap fast flow Q column (5 mL) and eluted against a linear salt gradient of 50-1000 mM NaCl at pH 7.6. Fractions containing synuclein were determined by SDS-PAGE and subjected to size exclusion chromatography using a Superdex 200 increase 10/300 GL column on an ÄKTA FPLC M system (Cytiva). The pure fractions were pooled and concentrated to ∼4 mg/ml. Protein was aliquoted and stored at -80 _°_C.

### Thioflavin T aggregation assays

The kinetics of fibrillogenesis by aSyn A53T and other variants were monitored according to 20 µM ThT (Sigma) fluorescence at 37 _°_C using a CLARIOstar microplate reader (BMG Labtech) with excitation at 450 nm and emission at 482 nm. The concentrated protein was diluted with buffer (25 mM Tris pH 7.6, 50 mM NaCl, plus different concentrations of calcium) to a final concentration of 1.5 mg/ml (∼100 µM). A final 150 µl reaction volume was placed in each well of a clear-bottom 96-well plate (Greiner) and sealed with transparent sealing film (Perkin Elmer). The plate was subjected to dual orbital shaking at 500 rpm and fluorescence signal was detected at 15 min intervals. Kinetic reactions were stopped when the steady-state phase of fibrils had formed. Seeds were generated using homogenized (vortexed) mature fibrillar material and then sonicated in a water bath at room temperature for 1 h. Aliquots of seeds were added to the top of final reaction volumes. Fibrillation kinetics were analyzed using Profit (https://www.quansoft.com/). Raw data were normalized to the final fluorescence signal by setting the first 10 time-points (150 minutes) and the last 10 time-points (150 minutes) as baselines. Lag time was attained by fitting a slope to the fibrillar sigmoid curve between 30 % and 70 %. Lag time was acquired by formulating the obtained slope and constant using the equation: *y* = *slope x* + *constant*, where *x* is the lag time^63^.

### Mass spectrometry of Ca^2+^-aSyn complexes

Samples of purified aSyn variants (4 mg/mL) in 25 mM Tris pH 7.6 and 50 mM NaCl—including wild-type, A53T, C-terminal-truncated (residues 1-109)—were diluted in 50% methanol and 50% double-distilled water (ddH_2_O) (v/v) to a final concentration of 2 µM. The 4 mg/ml aSyn sample plus 3.6 mM Ca^2+^ was prepared by adding a stock solution of 100 mM CaCl_2_ to purified protein and gently mixing with a pipette, before incubating at room temperature for 15 min. The sample was diluted with 50% methanol/50% ddH_2_O (v/v) to a final protein concentration of 2 µM. For all samples, 0.01% formic acid was added prior to undergoing MS analysis. For MS, samples were injected via the LockSpray Exact Mass Ionization Source (Waters) with a syringe pump (Harvard Apparatus), and held at a flow rate of 3 µL/min throughout the analysis. The mass of intact proteins was determined using a Waters Synapt G2 HDMS mass spectrometer (Waters). The deconvolution of ESI mass spectra of intact/reduced proteins was performed using the MaxEnt1 algorithm in MassLynx 4.1 software (Waters).

### NMR titration experiments

^15^N-labeled synuclein variants were generated according to protocols published previously^24^. In brief, overnight cultures were spun, collected, and resuspended in M9 solution containing 1 mg/ml ^15^NH_4_Cl and 2 mg/ml glucose. The procedures for overexpression and purification of ^15^N-labeled synuclein variants were identical to those described above for LB-cultured aSyn protein. All NMR experiments were performed at 15 _°_C on a Bruker AVANCE III 600 MHz spectrometer equipped with a TXI cryogenic probe. NMR samples were prepared in 25 mM HEPES pH 7.5, 50 mM NaCl, and 5 % D_2_O. All water-flipped ^15^N-edited HSQC spectra were acquired with 2048 (^1^H) and 256-300 (^15^N) time domain points, and 8-12 scans of 70-100 μM aSyn samples to avoid long-term data collection. Long-term spectral acquisition at high temperatures may induce unwanted protein aggregation during the acquisition process in the presence of calcium. All ^1^H-^15^N HSQC spectra were processed using NMRPipe^64^ and analyzed using NMRFAM-Sparky^65^.

A series of calcium titration NMR experiments were conducted to inspect how Ca^2+^ ions bound to aSyn A53T at the amino acid level. We conducted ^1^H-^15^N HSQC experiments using different concentrations of calcium (ranging from 0-20 mM) and a fixed aSyn A53T concentration at 70 μM. The titration data were used to confirm the calcium saturation concentration for aSyn A53T, in line with our ThT kinetics and PRE-NMR experiments. Calcium-induced peak shifts were further analyzed using a weighting function 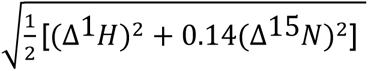 to calculate chemical shift perturbations (CSP) ^66^.

### Paramagnetic relaxation enhancement (PRE) NMR spectroscopy

As wild type aSyn lacks cysteines for conjugating PRE MTSL probes, we prepared the synuclein A53T G14C, S42C, and A140C mutant variants separately by means of site-directed mutagenesis. ^15^N-labeled aSyn A53T cysteine variants were generated as described above for other synuclein variants. The ^15^N-aSyn A53T cysteine variants were desalted using PD gravity desalting columns (Cytiva) to remove residual βME in the elution buffer, thereby ensuring that no disulfide bonds formed. The desalted sample was immediately conjugated to (1-oxy-2,2,5,5-tetra-methyl-3-pyrroline-3-methyl)-methanesulfonate (MTSL, Santa Cruz Biotechnology) by incubating it with a 10-fold molar excess of MTSL overnight at 4 _°_C. Excess MTSL was removed by means of desalting columns. Diamagnetic spectra were obtained by adding 10 mM βME in solution. The ^15^N-edited HSQC spectra of each cysteine mutant were collected under various experimental conditions (paramagnetic, diamagnetic, addition of 20 mM CaCl_2_) using an AVANCE III 600 MHz spectrometer (Bruker) at 15 _°_C. NMRFAM-Sparky was used to obtain the integrated peak heights that were used to calculate paramagnetic/diamagnetic intensity ratios.

### Transmission electron microscopy (TEM)

aSyn fibrils were initially cultivated at 1 mg/mL in buffer containing 25 mM Tris pH 7.6, 50 mM NaCl with or without 20 mM CaCl_2_. Reactions of 200 µL per tube were incubated at 37 _°_C with constant shaking and stopped after 2 weeks. The resulting aqueous fibril solution was sonicated in a water bath at room temperature to seed a second round of fibril preparation under the same cultivation conditions. A single iteration was achieved by adding 50 µl of seed to the 200 µl reaction volume in each tube.

To prepare negatively stained TEM samples, aqueous fibril solution was sonicated for 1 h in a water bath and then subjected to vortexing immediately prior to spreading the sample on glow-discharged 200 mesh Formvar/carbon-supported copper grids. The grids were then washed with ddH_2_O and stained twice with 2% uranyl acetate. Each step was allowed to proceed for 30-60 sec, before being gently dried with filter paper. Samples were imaged on a JEOL 1400 or an FEI Tecnai F20 TEM system in the cryo-EM facility of Academia Sinica, Taiwan.

### Cryogenic-electron microscopy data collection and processing

Mature aSyn fibrils were centrifuged at 15,000 rpm for 20 min to separate the remaining monomers. The pellet was resuspended by vortexing for 1 min in buffer (25 mM Tris pH 7.6, 50 mM NaCl). Samples were placed in a water bath and sonicated for 10 min, before being applied to glow-discharged Cu grids (Quantifoil R1.2/1.3, 300 mesh). The grids were blotted at 4 °C and 100% humidity, then blotted for 3 sec using a force setting of 10, before being vitrified via plunge-freezing in liquid ethane cooled by liquid nitrogen using an FEI Vitrobot Mark IV system. The cryo-EM data were acquired using a Titan Krios microscope (Thermo Scientific) operated at 300 kV with a magnification of 105,000X and equipped with a Gatan K3 camera and a BioQuantum energy filter with a slit width of 20 eV, resulting in a pixel size of 0.415 Å at super-resolution mode. Each micrograph was dose-fractionated to a total of 40 frames, and the electron dose rate was 41.64 e^−^/Å^2^. The defocus values were set at -1.2, -1.7 or - 2 µm for calcium-incubated aSyn A53T fibrils (aSyn A53T-Ca), and at -1, -1.4 or -1.8 µm for aSyn A53T fibrils without calcium (aSyn A53T). We collected 2663 and 1799 raw micrographs of aSyn A53T and aSyn A53T-Ca, respectively to reconstruct 3D maps of both fibril types. We used RELION 3.1^67, 68^ to conduct the entire cryo-EM process. Motion correction using a CPU-based RELION solution was applied to correct movies for drifting and for binning by a factor of 2 to yield a pixel resolution of 0.83 Å. Contrast transfer function (CTF) values were estimated in CTFFIND-4.1^69^. Helical reconstruction was performed carefully by following a protocol described previously^67, 70^.

We selected 13,968 filaments manually from 769 micrographs showing dispersed aSyn A53T-Ca fibrils. We extracted 167,252 fibril segments using a 280-pixel box at a 10% inter-box distance. 2D classification was carried out with a regularization value of T = 8. Segments assigned to appropriate 2D classes were selected to create initial 3D maps for crossover distance estimation. The initial helical rise at 4.75 Å was calculated from the 2D class layer line profile. Crossover of fibrils was roughly estimated from large boxed segments (500-800 pixels), before being confirmed using the helix_inimodel2d script in RELION 3.1. The initial helical twist at -1_°_ was calculated from the estimated crossover distance (850-900 Å). Three rounds of 3D classifications were conducted using T ranging from 8 to 20, resulting in 54,530 particles contributing to a reconstruction resolution of 3.63 Å. The best 3D-classified group was selected for initial 3D auto-refinement in RELION. Refinement by means of local searches using per-particle contrast transfer function (CTF) and Bayesian polishing resulted in final determined helical twist and rise values for aSyn A53T-Ca of 179.51_°_ and 2.375 Å, respectively. The aSyn A53T-Ca map was resolved to 2.7 Å using an estimated B factor of -75.06 Å^2^.

For the aSyn A53T fibrils, we selected 17,734 aSyn A53T fibrils, manually segmented them, and extracted them as boxes of 280 pixels with a 10% inter-box distance. We initially used 175,275 segments for 2D classification, with a regularization value of T = 6 or 8 to limit bad segments. Then, 109,125 segments were chosen to perform 3D classification (K=1) using an initial helical rise of 4.7 Å and an initial helical twist of - 1.5_°_, which were estimated from 2D averages of a large segment (800 pixels). A second 3D classification was carried out with parameters T=6, helical rise=2.375 Å, and helical twist=179.5_°_ to classify pseudosymmetric fibrils. One of the four 3D classes comprising 46,892 segments of sufficient quality was chosen for further refinements. Refinement with local searches of helical parameters achieved a 3.94 Å map, with a helical rise of 2.37 Å and a helical twist of 179.30_°_. The refined map was post-processed to a 3.42 Å map using a sharpening B-factor of -119 Å^2^. Data processing for both the aSyn-A53T and aSyn A53T-Ca maps is summarized in Table 1.

**Table 1.**
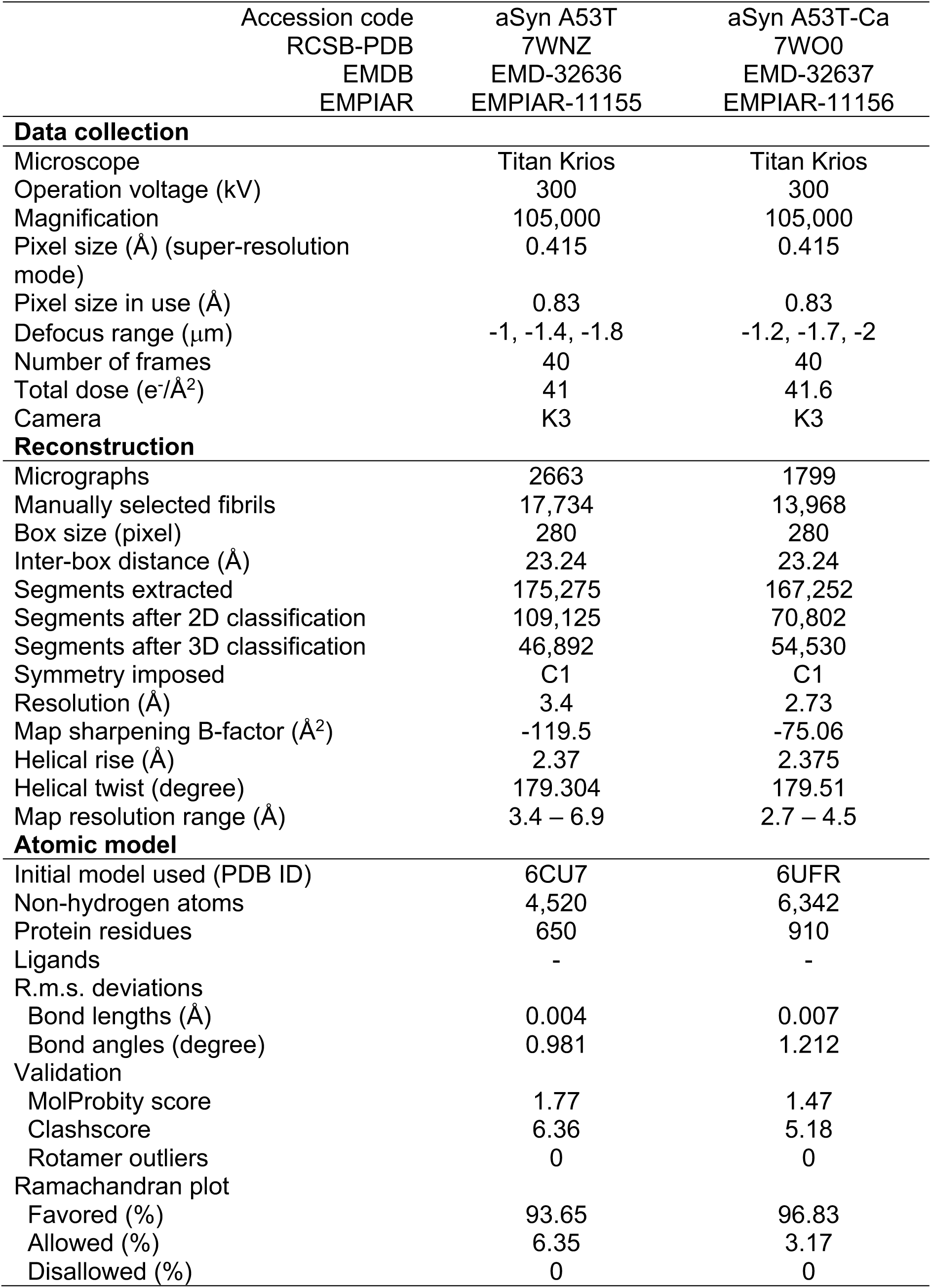
Cryo-EM and structural data for aSyn A53T and aSyn A53T-Ca.

### Building of aSyn fibril structures

Both of the aSyn A53T and A53T-Ca atomic models were built into RELION density maps using Coot 0.9.5^71^. The structures of wild type aSyn (PDB 6CU7^53^) and the aSyn E46K mutant variant (PDB 6UFR^54^) were used as initial templates for aSyn A53T and A53T-Ca, respectively, because of their similar structural folds. To avoid misleading model building of the polymorphic aSyn fibrils, we extracted a short 5-aa β-sheet from both models, docked it in a well-resolved β-sheet density in the map, and then mutated it to a penta-alanine β-sheet. A single chain was initially built by means of alanine extension in the N- and C-terminal directions. Checkpoint residues (including Y39, H50, V52, V74, V77, I88, and F94) were used to accurately assign sidechains for the whole single chain model in Coot. The built chain was then copied and docked repeatedly into the map to generate separate initial models of aSyn A53T and A53T-Ca. Phenix 1.18 real-space refinement^72^ was utilized to optimize the sidechain rotamers, bond angle, bond length, dihedral angles, and inter-chain distances of initial models. Refined structures and manually corrected structures were iteratively inspected in Coot to achieve good model-to-map cross-correlation and structural quality. Finalized structures of aSyn A53T and A53T-Ca were validated using MolProbity^73^ and their characteristics are summarized in Table 1.

## Data availability

The cryo-EM density maps of aSyn A53T and aSyn A53T-Ca have been deposited in the Electron Microscopy Data Bank under accessions EMD-32636 and EMD-32637, respectively. Motion corrected micrographs of aSyn A53T and aSyn A53T-Ca were deposited in Electron Microscopy Public Image Archive (EMPIAR) under accessions EMPIAR-11155 and EMPIAR-11156. The accession numbers of associated structure coordinates of aSyn A53T and aSyn A53T-Ca deposited in the RCSB Protein Data Bank are 7WNZ and 7WO0, respectively.

## Acknowledgments

K.-P.W. is supported by an Academia Sinica (AS) Career Development Award (AS-CDA-110-L03) and by the National Council of Science and Technology (NSTC-111-2113-M-001-026-MY2). We appreciate Dr. Meng-Ru Ho from the Biophysics Instrumentation Laboratory, Institute of Biological Chemistry, AS, and Dr. Chris Shu-Chuan Jao from the Biophysics Core Facility (BCF), AS, for their assistance and valuable discussions on aSyn biophysical characterization. We thank AS High-field NMR Center (HFNMRC), AS Common Mass Spectrometry Facilities (ASCMSF), and the AS Cryo-EM facility (ASCEM) for their technical support. BCF, HFNMRC, ASCMSF, and ASCEM are supported by Academia Core Facility and Innovative Instrument Projects (AS-CFII-111-201, AS-CFII-111-209AS-CFII-112-103, AS-CFII-112-210).

## Notes

### Competing Interest Statement

The authors have declared no competing interest.

