## supporting information for "Calcium modulates intramolecular long-range contacts to form a polymorphic α-synuclein A53T fibril"

This PDF file contains 4 supplementary figures in 6 pages.

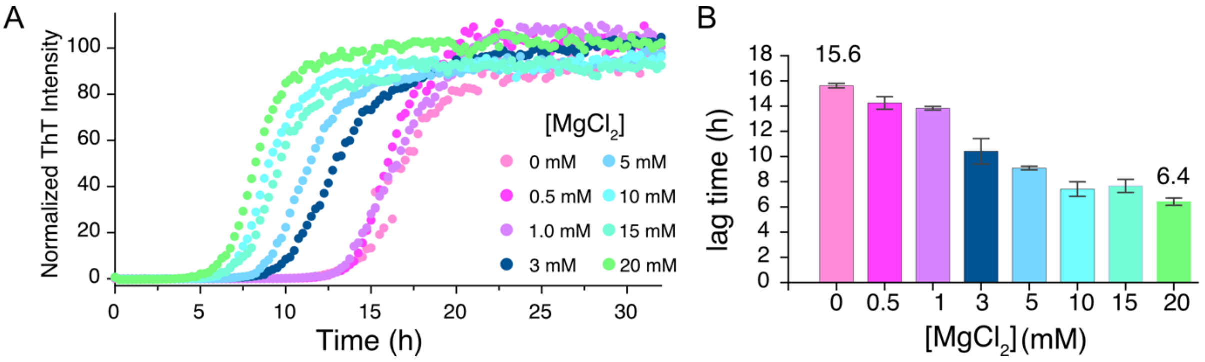


**Supplementary Figure 1. Impacts of $MgCl_2$ on the ThT aggregation kinetics of aSyn A53T.**

(A). The aggregation kinetics of aSyn A53T were accelerated by increasing concentrations of $MgCl_2$. (B). A lag time analysis indicates that aggregation rates were maximized under conditions of 10-20 mM $MgCl_2$.

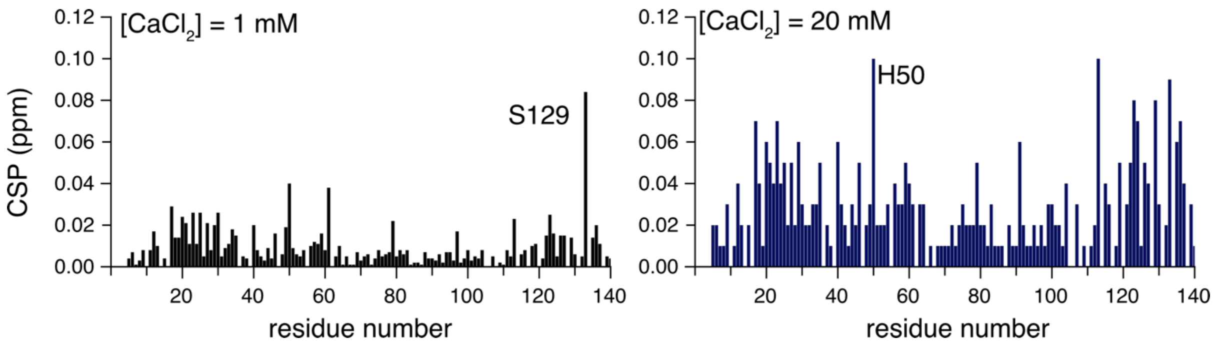


**Supplementary Figure 2. Chemical shift perturbations (CSP) of aSyn A53T in the presence of high [$CaCl_2$].**

aSyn A53T (70 µM) is mildly influenced by 1 mM $CaCl_2$ (left), with the majority of CSP values being below 0.02 ppm. S129 proved to be the most sensitive residue upon $CaCl_2$ titration. However, in the presence of 20 mM $CaCl_2$, high CSP values are apparent across the entire sequence, with residues 20-35, 120-140, 58-62, and 74-80 being greatly perturbed.

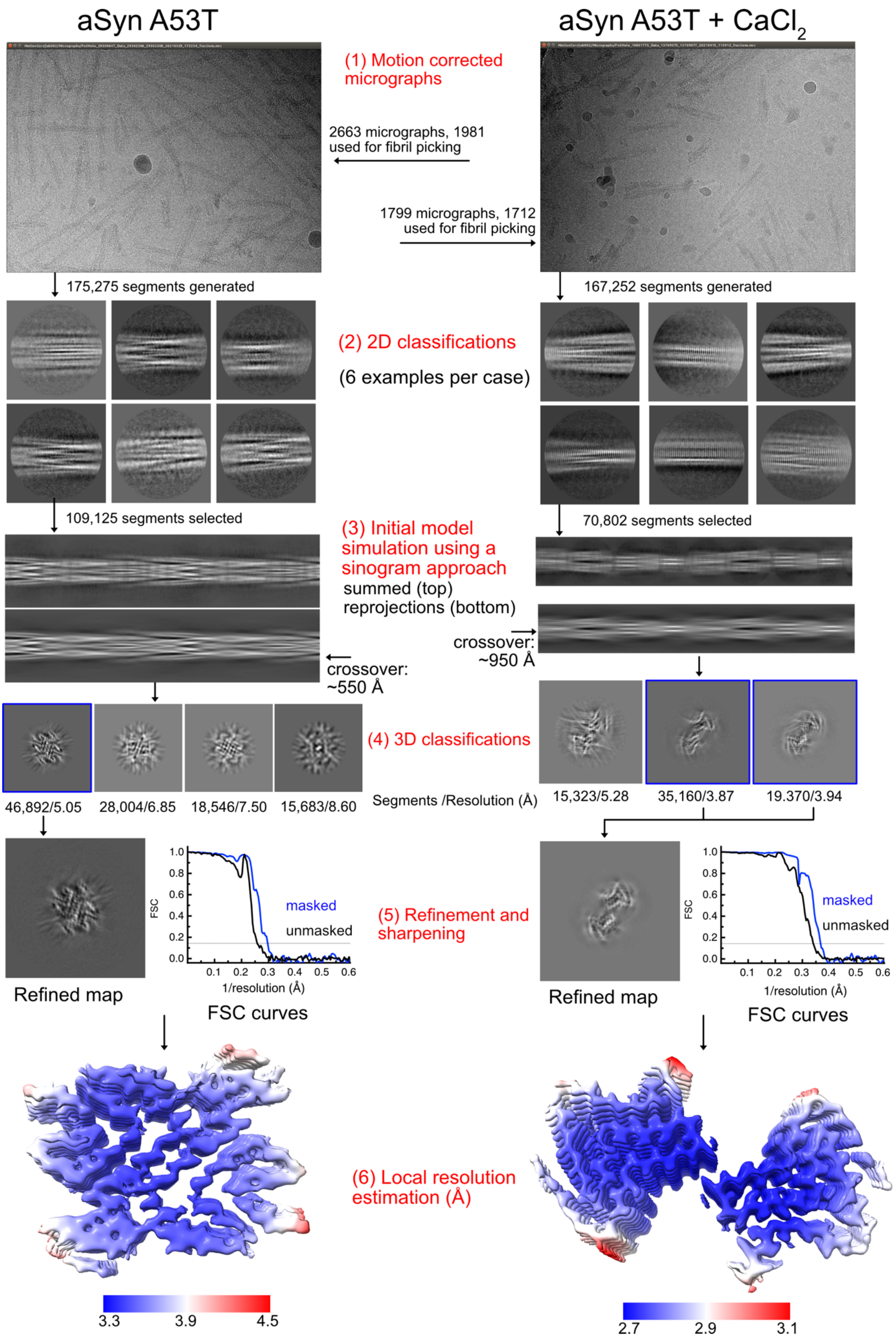
aSyn A53T
aSyn A53T + CaCl2
(1) Motion corrected micrographs
2663 micrographs, 1981 used for fibril picking
1799 micrographs, 1712 used for fibril picking
175,275 segments generated
167,252 segments generated
(2) 2D classifications
(6 examples per case)
109,125 segments selected
70,802 segments selected
(3) Initial model simulation using a sinogram approach
summed (top)
reprojections (bottom)
crossover: ~550 Å
crossover: ~950 Å
(4) 3D classifications
46,892/5.05
28,004/6.85
18,546/7.50
15,683/8.60
Segments /Resolution (Å)
15,323/5.28
35,160/3.87
19.370/3.94
(5) Refinement and sharpening
Refined map
FSC curves
FSC
masked
unmasked
1/resolution (Å)
(6) Local resolution estimation (Å)
3.3
3.9
4.5
2.7
2.9
3.1

**Supplementary Figure 3. Cryo-EM data processing of aSyn A53T and aSyn A53T-Ca fibrils.**
The workflow pipelines for aSyn A53T (left) and aSyn A53T-Ca (right) are summarized. There were six major steps: (1) motion corrections of micrographs and manual selection of fibrils; (2) 2D classification; (3) initial model generation; (4) 3D classification; (5) 3D refinement and post-processing; and (6) local resolution estimation. Key numbers associated with each step are presented. For the 3D classification step, good groups (highlighted by blue boxes) were selected for further 3D refinement calculation. Local resolution was estimated in RELION 3.1 and the different resolution ranges of aSyn A53T (3.3-4.5 Å) and aSyn A53T-Ca (2.7-3.1 Å) are illustrated.

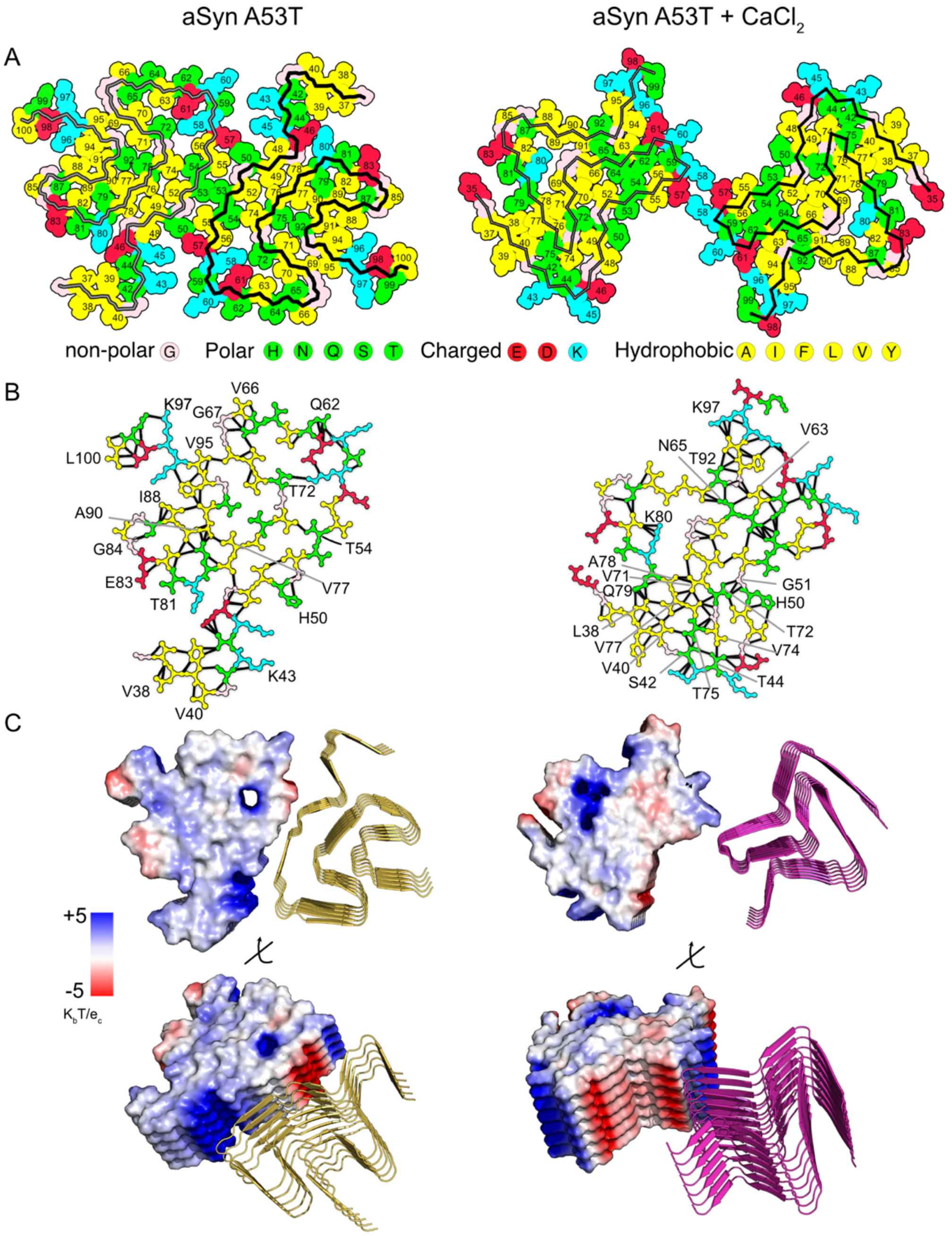


**Supplementary Figure 4. The intrachain and interchain interactions of aSyn A53T and aSyn A53T-Ca fibrils.**

(A). Sidechain occupations for both aSyn fibril types are presented, in which the different colors code for polar, non-polar, hydrophobic or charged residues, as per the legend. aSyn A53T assembles via a steric zipping mechanism, whereas aSyn A53T-Ca protofilaments pair via charge-charge interactions. (B). Intrachain van der Waal's interactions (as determined in ChimeraX 1.3) are linked by black lines, with key residues involved in hydrophobic contacts indicated. The color codes are identical to those in (A). (C). Electrostatic surface potentials of both aSyn fibril types, as generated using PyMOL 2.5.0. Blue and red colors represent positive and negative charges, respectively, within a range of +5 to -5 $K_bT/e_c$. The charge distributions reveal a larger hydrophobic patch (white) for aSyn A53T-Ca (right) relative to aSyn A53T (left). Sideview of the protofilament interfaces revealing that strong electrostatic potentials are one of the factors promoting protofilament assembly.
